# Cas9-Assisted Targeting of CHromosome segments (CATCH) for targeted nanopore sequencing and optical genome mapping

**DOI:** 10.1101/110163

**Authors:** Tslil Gabrieli, Hila Sharim, Yael Michaeli, Yuval Ebenstein

**Author notes:** Correspondence may also be addressed to Yael Michaeli.

## Abstract

Variations in the genetic code, from single point mutations to large structural or copy number alterations, influence susceptibility, onset, and progression of genetic diseases and tumor transformation. Next-generation sequencing analysis is unable to reliably capture aberrations larger than the typical sequencing read length of several hundred bases. Long-read, single-molecule sequencing methods such as SMRT and nanopore sequencing can address larger variations, but require costly whole genome analysis. Here we describe a method for isolation and enrichment of a large genomic region of interest for targeted analysis based on Cas9 excision of two sites flanking the target region and isolation of the excised DNA segment by pulsed field gel electrophoresis. The isolated target remains intact and is ideally suited for optical genome mapping and long-read sequencing at high coverage. In addition, analysis is performed directly on native genomic DNA that retains genetic and epigenetic composition without amplification bias. This method enables detection of mutations and structural variants as well as detailed analysis by generation of hybrid scaffolds composed of optical maps and sequencing data at a fraction of the cost of whole genome sequencing.

## Introduction

Since the introduction of next generation sequencing (NGS), tremendous progress has been achieved in all fields of biology. The declining costs and growing availability of NGS have made it the method of choice for genetic analysis and related applications. Nevertheless, NGS suffers from several limitations that prevent extraction of the full extent of information associated with the genome such as data on repetitive elements and large structural variations (SVs) (1). The major source of these limitations is that NGS technology relies on the alignment of millions of short reads, of up to several hundred bases, to produce a final genome sequence; therefore, long-range information is lost. Consequently, the reference human sequence contains gaps mainly due to structural and copy number variations that could not be resolved by sequencing. Furthermore, different experimental methods and computational analysis of NGS datasets applied to the same human DNA samples show disappointingly low levels of overlap (2). Recently developed single-molecule sequencing techniques such as single-molecule real-time sequencing (SMRT) and nanopore sequencing provide access to larger variations due to typical read lengths of several thousands of bases (3, 4); however, whole genome assemblies using these techniques is challenging. Although in many cases only a specific locus is of interest, whole genome analysis must be performed due to the absence of efficient enrichment techniques, incurring high costs and resulting in complex data analysis. Current enrichment schemes are based on PCR amplification or hybridization capture of specific regions. Both PCR amplification and non-uniform capture efficiency are inherent sources of bias with these approaches, especially in highly repetitive areas. Additionally, both methods require prior knowledge of the sequence, which limits them to well characterized, non-variable areas. Targeted analysis of genomic regions longer than 200 kilo-base pairs (bp) is extremely complicated, and therefore targeting schemes usually focus on exons and neglect the introns and regulatory regions (5, 6).

Here, we demonstrate the use of Cas9-Assisted Targeting of CHromosome segments (CATCH) for the enrichment of large genomic fragments for genetic analysis. The target DNA is extracted as intact molecules enabling multiscale analysis of short and long-range information. We use a Cas9 nuclease that is guided to its target cleavage site by sequence homology of an RNA subunit. Our method is based on capturing cells in gel plugs for lysis and extraction of gel-trapped high-molecular weight chromosomal DNA. This is followed by *in vitro* digestion of the DNA with the Cas9 enzyme at two sites flanking the locus of interest (Figure 1). We have previously shown that CATCH can be used for efficient cloning of large genomic segments via Gibson assembly (7, 8). In the current study we show that the enriched DNA can be analyzed in detail at various genomic scales at reduced complexity and cost relative to whole genome sequencing methods.

**Figure 1.**
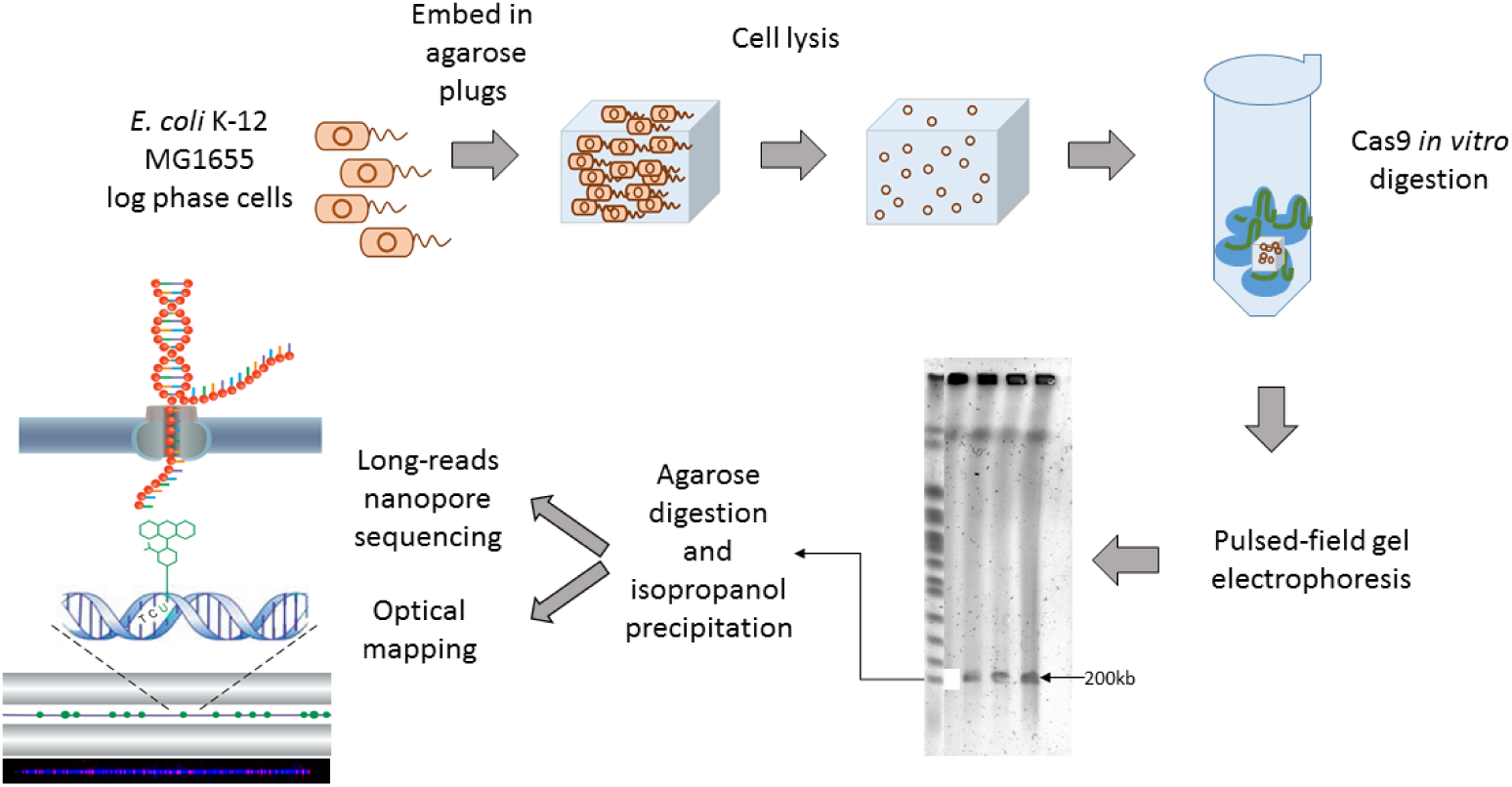
Schematic of CATCH technique. Log-phase *E. coli* cells were embedded in an agarose gel and lysed. Genomic DNA was digested using guided Cas9, and the target DNA was separated by PFGE. The desired band was excised from the gel, and the DNA was isolated. The target DNA was analyzed by nanopore sequencing and optical mapping.

For single base-pair resolution analysis, we use the MinION nanopore sequencing platform (Oxford Nanopore Technologies, Figure 1). The MinION is a small portable device that can be connected to a personal computer. Sequencing is based on threading a DNA molecule through a nanoscale protein pore while measuring current across the pore. The DNA bases cause a current modulation such that each base entering the pore is characterized by a different current level, thus identifying the base in each position. This technology enables long reads with typical lengths of several kbp and up to 300 kbp (9), which makes it suitable for assembly of repetitive regions and medium-size structural variations.

For characterizing larger SVs, ranging up to hundreds of kbps, we use a technique called optical genome mapping (10–12). This technology, commercialized by Bionano Genomics Inc., is based on electro-kinetically driving DNA molecules through an array of nanochannels that confine the DNA in an extended state. DNA is labeled with fluorescent molecules at specific sequence motifs by an enzymatic reaction, and labels are imaged by a fluorescence microscope as a linear fluorescent barcode that reports on the genomic loci of the molecules. The pattern of fluorescent spots along the DNA is aligned to an *in silico* reference map or assembled *de novo* to create consensus contigs spanning many mega base pairs (Mbp). This technology permits visualization of multiple DNA molecules up to several Mbp long and allows high-throughput detection of complex structural features and the identification of structural variants (13). Combining the two data sets by aligning local sequence information to the optical map can resolve complex genomic regions and close gaps in the genome sequence (14). As a proof of concept for the developed technique, we isolated a 200 kbp fragment from the *Escherichia coli (E. coli)* genome and analyzed the fragment by optical mapping and nanopore sequencing analysis.

## MATERIAL AND METHODS

### Guide RNA design and preparation

Two guide RNAs were designed in order to extract a ~200 kbp fragment from the bacterial genome of *E. coli* str. K-12 substr. MG1655. Guide RNAs were designed as previously described (see supplementary information, (7)). Briefly, the online CRISPR design tool (http://crispr.mit.edu)was used to locate a 20 bp recognition sequence followed by a NGG protospacer-adjacent motif (PAM) with minimal identity to other parts of the genome. A 117 bp template DNA that contains T7 promoter, recognition sequence, crRNA, and tracrRNA was generated from an overlap PCR of three primers. In order to create the gRNA, an *in vitro* transcription (IVT) reaction was performed on the template DNA with HiScribe T7 RNA synthesis kit (NEB) according to manufacturer’s instructions. gRNA was purified by phenol-chloroform purification and isopropanol precipitation. RNA was resuspended in RNase-free water and stored at −80 °C until use.

### Plug preparation

*E. coli* K-12 MG1655 cells were grown in LB, and log phase cells were embedded in low melting agarose gel plugs at a concentration of ~2 x 10^9^ cells per ml. Cells were resuspended in cell suspension buffer (CHEF mammalian DNA extraction kit, Bio-Rad) prior to plug casting. Plugs were treated with lysozyme (Sigma) followed by proteinase K (Qiagen) treatment in lysis buffer (BioNanoGenomics). RNase digestion (Qiagen) was then performed in TE buffer. After washing, plugs were stored at 4 °C until use.

### In-gel Cas9 digestion

gRNA and Cas9 were pre-assembled prior to the fragmentation reaction. Plugs were washed with 10 mM Tris-HCl (pH 8) and then with Cas9 reaction buffer (NEB). Plugs were then cut into three equal parts (~30 µl each), and each part was incubated with preassembled gRNA-Cas9 (NEB). Plugs were then incubated with Proteinase K in order to remove excess Cas9 bound to the DNA. Protocols are detailed in the supplementary information.

### Separation by pulsed field gel electrophoresis (PFGE)

A 0.9% solution of sea plaque low melting agarose gel (Lonza) was prepared in 0.3X TBE. 15 digested plugs were loaded onto the gel and run on a Rotaphor instrument (Biometra) for 24 hours in 0.25X TBE buffer. The band corresponding to ~200 kbp was cut from the gel, and DNA was extracted from the agarose using beta-agarase (Thermo Fisher Scientific) and isopropanol precipitated. Details are given in the supplementary information.

### Nanopore sequencing

For construction of the nanopore sequencing library, a low input expansion pack kit in combination with SQK-MAP007, R9 version kit (Oxford Nanopore Technologies) were used according to the manufacturer’s instructions with minor modifications (see supplementary information for details). Libraries were loaded on R9.4 flowcells, and the MinION sequencing device was run for 48 hours. Reads were generated by the MinKNOW control software and base-called using the Metrichor software. Fast5 files that either passed (’pass reads’) or failed (’fail reads’) base-calling quality metrics were converted to FASTA or FASTQ files using poretools (version 0.6.0) with–type best option (15). Alignment of reads to the *E. coli* reference genome was performed using BWA-MEM (version 0.7.12 (16)) with -x ont2D parameters, and coverage was calculated using the Galaxy wrapper for BEDTools genomecov (version 2.26.0 (17, 18)).*De novo* assembly was performed on 2D pass reads using Canu (version 1.4, (19)) with default parameters, specifying a genome size of 200 kbp. Long reads were converted into a CMAP file imitating an optical map using the Knickers software (version 1.5.5) provided by BioNano Genomics by specifying the Nt.BspQI recognition sequence. The bam2R function from the R package deep SNV was used to evaluate the error rate of the reads.

### Optical mapping: Sample preparation, imaging, and analysis

The nicking endonuclease Nt.BspQI (NEB) was used to introduce single-stranded nicks into the double-stranded DNA (dsDNA) at specific sequences (GCTCTTCN^). Taq DNA polymerase (NEB) that was supplemented with dUTP-atto 532 (Jena Bioscience) in addition to dATP, dGTP, and dCTP (Sigma) was used to label the DNA at the nicking sites. Nicks were then ligated with Taq DNA Ligase (NEB) and nicked-labeled DNA was stained with yoyo-1 (DNA stain, BioNanoGenomice). Loading of DNA in nanochannels and imaging were performed using an Irys instrument (BioNano Genomics). Detection of imaged molecules and fluorescent labels along each molecule was performed by AutoDetect (version 2.1.4, BioNano Genomics). Alignment to the reference genome and *de novo* assembly were performed using IrysView software (version 2.3, BioNano Genomics). Details of experimental protocols are given in supplementary information.

## RESULTS

### Fragmentation and isolation of a large genomic target sequence using CATCH

Our method for enrichment of a large genomic locus is based on *in vitro* fragmentation with S. *pyogenes* CRISPR/Cas9. As a proof of concept we used the *E. coli* strain K-12 MG1655, which has a genome size of ~4.5 Mbp. In order to cut the bacterial genome at two positions, we designed two gRNAs with recognition sequences flanking the ~200kbp target fragment (see supplementary information and (7)). High molecular weight DNA was extracted by lysis of bacterial cells in gel plugs; this strategy minimizes mechanical shearing and maintains DNA integrity. The gel plugs were soaked with Cas9 and gRNAs to excise the target region, and DNA was loaded on a low melting agarose gel for separation by pulse-field gel electrophoresis (PFGE). In this experiment, 15 plugs with a total of approximately 12 µg of DNA were loaded onto the gel, which should theoretically yield about 500 ng of the 200 kbp fragment. A band corresponding of approximately 200 kbp was clearly observed in the gel (Figure 1). This band was cut out of the gel, and the agarose was enzymatically digested. Following purification, a total of 200 ng target DNA was recovered from the gel, which corresponds to 40% recovery. Of this, 125 ng were used for nanopore library construction and the remaining 75 ng were used for optical mapping.

### Nanopore sequencing analysis of targeted genomic fragment

Following isolation of the target DNA by CATCH, we used long-read nanopore sequencing to characterize our sample and estimate the level of enrichment. After mechanical shearing according to the manufacturer’s protocol and preparation of the sequencing library from the fragmented sample, we obtained 11,607 reads in a single experiment, with 1,425 reads that were considered “pass reads”. Of the pass reads, 99.6% were aligned to the *E. coli* genome, and half (~49%) of pass reads were aligned to the 200 kbp target (Figure 2, Figure S1). This fragment corresponds to less than 5% of the bacterial genome, which implies a 10-fold enrichment of the target. When both pass and fail reads were analyzed, 50% of the reads were aligned to the *E. coli* genome. This could be a result of the higher error rate when 2D base-calling fails (Figure S2). Nevertheless, approximately 46% of these reads were aligned to the 200 kbp fragment, despite the fact that this target region corresponds to less than 5% of the bacterial genome. These reads covered the entire target region at an average sequencing depth of 74x, while the average sequencing depth of the rest of the genome was 3.4x (Figure 2), indicating an enrichment factor (EF) of 21.7 (20). Pass reads were used for *de novo* assembly and yielded a single contig spanning the entire 200 kbp region with 99% identity to the reference. A minimal set of 17 reads were sufficient to generate complete coverage of the entire 200 kbp target region with the longest read covering 42,926 bp of the region (Figure S1).

**Figure 2.**
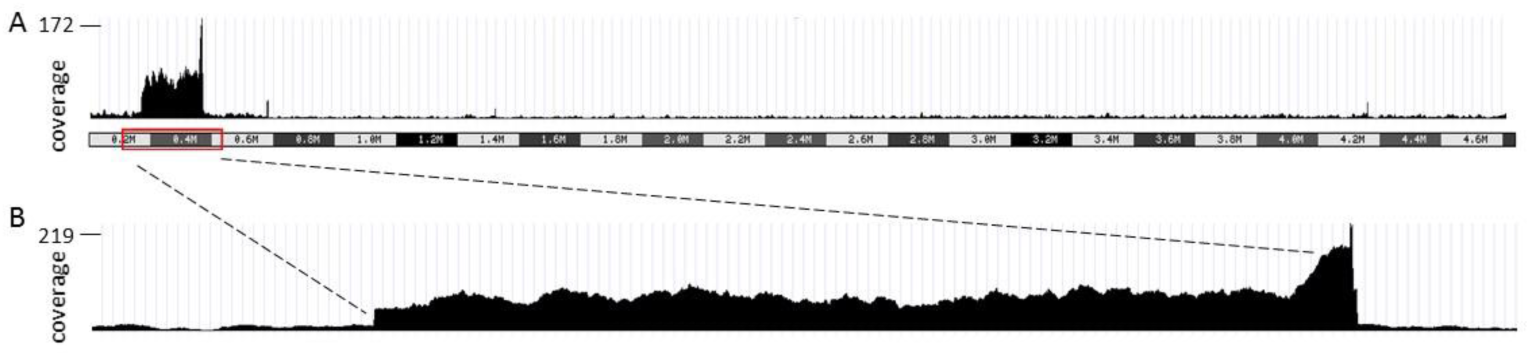
Coverage analysis of the enriched 200 kbp target region analyzed by nanopore sequencing. A. Read coverage along the entire *E. coli* genome. B. Read coverage of the 200 kbp target region. Results were visualized using the UCSC genome browser.

### Optical mapping analysis of targeted genomic region

Next, we used optical mapping to analyze the enriched DNA. This method is used when long range information is required regarding the structure of a target region. Mapping requires only knowledge of the Cas9 cleavage site regions and gives access to information such as large structural and copy number variation within the target region. We used nick translation to introduce a fluorophore at specific sequence motifs, in order to create a fluorescent molecular barcode. The labeled DNA was then stretched by forcing it into nano-channels using an electric field, allowing fluorescent imaging of its whole contour. Imaging was performed using an Irys instrument and IrysChip (BioNanoGenomics) composed of silicon fabricated nanochannel arrays with ~45-nm-wide channels, which forces the DNA to stretch uniformly. The detected molecules were then aligned to the reference genome. A total of 1,553 molecules above 75 kbp in length were aligned to the *E. coli* genome, with 1,405 molecules aligned to the 200 kbp fragment (~90%), out of which 76 spanned over 95% of the target region. The mean coverage across the target was 516 while the coverage across the rest of the genome was 17, indicating an enrichment factor of 209. In addition, unsupervised *de novo* assembly yielded three contigs, including one consensus contig that contained the entire 200 kbp target sequence and two other flanking contigs around the target sequence (Figure S3).

**Figure 3.**
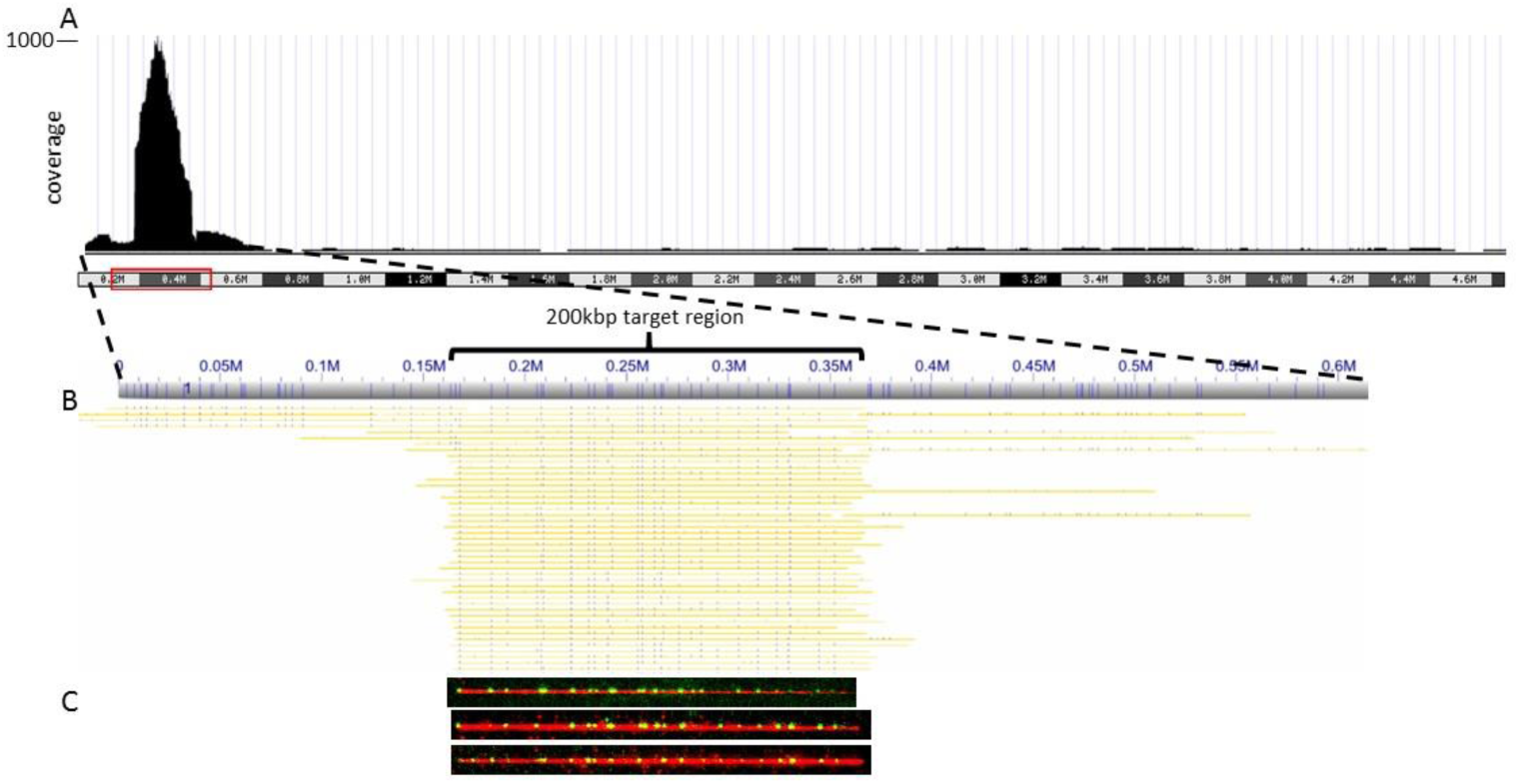
Coverage analysis of the enriched 200 kbp target region analyzed by optical mapping. A. Coverage of molecules longer than 50 kbp along the entire *E. coli* genome. Results were visualized using UCSC genome browser. B. Alignment of molecules (represented in yellow) to the *E. coli* reference (gray). Blue lines indicate Nt.BspQI nicking sites. C. Images of labeled molecules from the target region detected in the nanochannels.

### Hybrid scaffolding of nanopore sequencing reads and optical mapping data

One of the potential uses of the CATCH technique is the isolation of large genomic fragments that are poorly characterized and composed of extremely variable regions or repetitive areas. In such cases, the reference sequence poorly represents the actual structure of the region or is not available at all. Optical maps are generated from intact long molecules with a distinct pattern of known sequence motifs highlighted by fluorescence labeling. Long sequencing reads containing several of these sequence motifs can be directly aligned to the optical consensus map based on the arrangement of the motifs and without prior knowledge of the reference sequence. The hybrid scaffold is generated by anchoring the contigs generated from the sequencing data to the consensus map achieved by optical mapping thus providing sequence information on the regions between optical labels in the map (21). An example is shown in Figure 4. First, a consensus optical map is generated by assembling the individual molecular optical maps (light blue track). Then, nanopore reads are mapped to the optical consensus pattern of the nicking enzyme recognition sequences based on the pattern of motifs in the sequence read. Consensus sequence contigs are then generated from the overlapping mapped sequencing reads. This procedure can potentially increase the N50 of the sequence contigs and reveal large structural and copy number variations, which could not be characterized by other methods.

**Figure 4.**
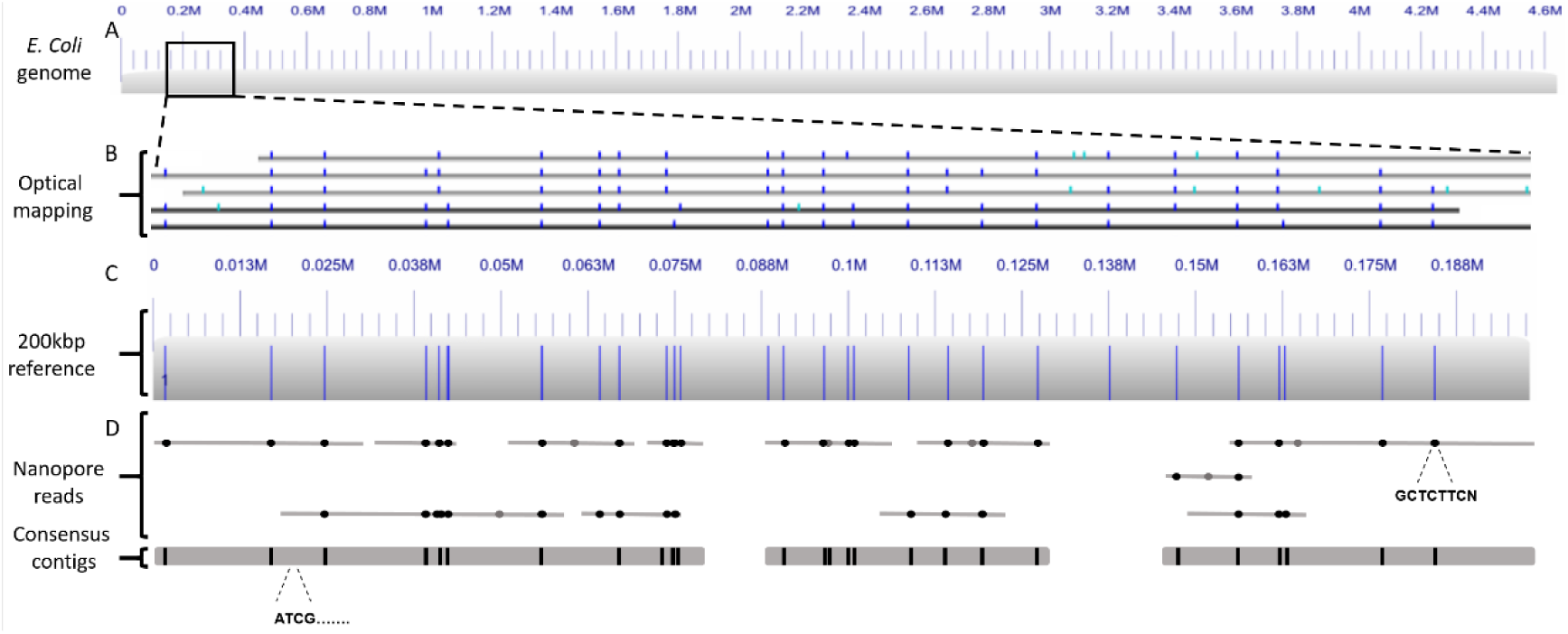
Generation of nanopore sequencing and optical mapping hybrid maps. A. Digitized representation of the *E. coli* genome reference. B. Digitized representation of molecules aligned to the 200 kbp target from optical mapping. Blue lines indicate detected Nt.BspQI nicking sites. C. Reference of the 200 kbp target region. Blue lines indicate expected Nt.BspQI nicking sites. D. Optical representation of long nanopore reads aligned to the 200 kbp reference and consensus contigs generated from overlapping reads. Nt. BspQI recognition sites are indicated by black dots. False positive Nt. BspQI sites are indicated by gray dots.

## DISCUSSION

Despite great progress in recent years, DNA sequencing techniques have several limitations in characterizing highly repetitive or highly variable areas. Additionally, the read depth required for accurate analysis of such areas is extremely high, which is cost prohibitive on a whole-genome scale. An outstanding challenge is to be able to profile intact, highly variable genomic regions spanning hundreds of thousands to several millions of bases at very high coverage. This includes long disease-associated genes or pathogenic macrosatellite arrays, where a full, multi-scale genetic analysis is needed. It has been shown that mutations not just in exons, but also in non-coding regions, are significantly correlated with disease class and severity.

Additionally, epigenetic markers are lost during amplification but also influence the state of a disease. Examples of such genomic areas are the *D4Z4* repeat array, which is related to Facioscapulohumeral muscular dystrophy (FSHD) (22), and the breast cancer-related genes *BRCA1* and *BRCA2* (23); these genes extend between 80 and 250 kbp and are known to have various pathogenic point mutations and structural rearrangements (24, 25). To facilitate early diagnosis of malignant transformation, genomic mutations must be detected when only a small population of cells is transformed. This requires sequencing depth (coverage) of 2-3 orders of magnitude deeper than a conventional whole genome sequencing run and an order of magnitude deeper than whole exome sequencing runs. Enrichment of target genomic loci using CATCH may enable comprehensive multi-scale analysis at reduced costs.

CATCH can provide the long-range information that is missing in other techniques by enabling isolation of intact high molecular weight DNA. As a proof of concept, we successfully excised and enriched bacterial DNA of an approximately 200 kbp fragment. We performed long-read nanopore sequencing to obtain sequence information at single base-pair resolution and optical mapping for large-scale structure analysis. In addition, we demonstrated that these two data types could be combined by hybrid scaffolding for elucidating sequence and structure. This will be useful in cases where a reference map is unreliable or non-existent. We envision that extending this technique to larger and more complex genomes will allow cost-effective access to data on a plethora of variable and poorly characterized genomic regions. Direct applications include analyzing highly repetitive plant genomes such as wheat (26) and in medical genomics focused on specific large pathogenic regions such as the *BRCA* (23) and *APC* (27) gene regions.

## REFERENCES

1. Treangen, T.J. and Salzberg, S.L. (2011) Repetitive DNA and next-generation sequencing:computational challenges and solutions. Nat. Rev. Genet., 13, 36–46.

2. Duan, J., Zhang, J.-G., Deng, H.-W. and Wang, Y.-P. (2013) Comparative studies of copynumber variation detection methods for next-generation sequencing technologies. PLoS One, 8, e59128.

3. Norris, A.L., Workman, R.E., Fan, Y., Eshleman, J.R. and Timp, W. (2016) Nanopore sequencing detects structural variants in cancer. Cancer Biol. Ther., 17, 246–53.

4. Huddleston, J., Chaisson, M.J., Meltz Steinberg, K., Warren, W., Hoekzema, K., Gordon, D.S., Graves-Lindsay, T.A., Munson, K.M., Kronenberg, Z.N., Vives, L., et al. (2016) Discovery and genotyping of structural variation from long-read haploid genome sequence data. Genome Res., 10.1101/gr.214007.116.

5. Bodi, K., Perera, A.G., Adams, P.S., Bintzler, D., Dewar, K., Grove, D.S., Kieleczawa, J., Lyons, R.H., Neubert, T.A., Noll, A.C., et al. (2013) Comparison of commercially available target enrichment methods for next-generation sequencing. J. Biomol. Tech., 24, 73–86.

6. Mamanova, L., Coffey, A.J., Scott, C.E., Kozarewa, I., Turner, E.H., Kumar, A., Howard, E., Shendure, J. and Turner, D.J. (2010) Target-enrichment strategies for next-generation sequencing. Nat. Methods, 7, 111–118.

7. Jiang, W., Zhao, X., Gabrieli, T., Lou, C., Ebenstein, Y. and Zhu, T.F. (2015) Cas9-Assisted Targeting of CHromosome segments CATCH enables one-step targeted cloning of large gene clusters. Nat. Commun., 6, 8101.

8. Jiang, W. and Zhu, T.F. (2016) Targeted isolation and cloning of 100-kb microbial genomic sequences by Cas9-assisted targeting of chromosome segments. Nat. Protoc., 11, 960–975.

9. Ip, C.L.C., Loose, M., Tyson, J.R., de Cesare, M., Brown, B.L., Jain, M., Leggett, R.M., Eccles, D.A., Zalunin, V., Urban, J.M., et al. (2015) MinION Analysis and Reference Consortium: Phase 1 data release and analysis. F1000Research, 4, 1075.

10. Samad, A., Huff, E.F., Cai, W. and Schwartz, D.C. (1995) Optical mapping: a novel, singlemolecule approach to genomic analysis. Genome Res., 5, 1–4.

11. Howe, K. and Wood, J.M. (2015) Using optical mapping data for the improvement of vertebrate genome assemblies. Gigascience, 4, 10.

12. Levy-Sakin, M. and Ebenstein, Y. (2013) Beyond sequencing: optical mapping of DNA inthe age of nanotechnology and nanoscopy. Curr. Opin. Biotechnol., 24, 690–8.

13. Cao, H., Hastie, A.R., Cao, D., Lam, E.T., Sun, Y., Huang, H., Liu, X., Lin, L., Andrews, W., Chan, S., et al. (2014) Rapid detection of structural variation in a human genome using nanochannel-based genome mapping technology. Gigascience, 3, 34.

14. Mostovoy, Y., Levy-Sakin, M., Lam, J., Lam, E.T., Hastie, A.R., Marks, P., Lee, J., Chu, C., Lin, C., Dzžakula, ZŽ., et al. (2016) A hybrid approach for de novo human genome sequence assembly and phasing. Nat. Methods, 13, 587–590.

15. Loman, N.J. and Quinlan, A.R. (2014) Poretools: a toolkit for analyzing nanopore sequence data. Bioinformatics, 30, 3399–3401.

16. Li, H. and Durbin, R. (2009) Fast and accurate short read alignment with Burrows-Wheeler transform. Bioinformatics, 25, 1754–1760.

17. Goecks, J., Nekrutenko, A., Taylor, J. and Galaxy Team, T. (2010) Galaxy: a comprehensive approach for supporting accessible, reproducible, and transparent computational research in the life sciences. Genome Biol., 11, R86.

18. Quinlan, A.R. and Hall, I.M. (2010) BEDTools: a flexible suite of utilities for comparing genomic features. Bioinformatics, 26, 841–842.

19. Koren, S., Walenz, B.P., Berlin, K., Miller, J.R. and Phillippy, A.M. (2016) Canu: scalable and accurate long-read assembly via adaptive k-mer weighting and repeat separation. bioRxiv.

20. Mertes, F., Elsharawy, A., Sauer, S., van Helvoort, J.M.L.M., van der Zaag, P.J., Franke, A., Nilsson, M., Lehrach, H. and Brookes, A.J. (2011) Targeted enrichment of genomic DNA regions for next-generation sequencing. Brief. Funct. Genomics, 10, 374–86.

21. Mostovoy, Y., Levy-Sakin, M., Lam, J., Lam, E.T., Hastie, A.R., Marks, P., Lee, J., Chu, C., Lin, C., Dzžakula, ZŽ., et al. (2016) A hybrid approach for de novo human genome sequence assembly and phasing. Nat. Methods, 13, 587–590.

22. Cabianca, D.S. and Gabellini, D. (2010) The cell biology of disease: FSHD: copy number variations on the theme of muscular dystrophy. J. Cell Biol., 191,1049–60.

23. Sluiter, M.D. and van Rensburg, E.J. (2011) Large genomic rearrangements of the BRCA1 and BRCA2 genes: review of the literature and report of a novel BRCA1 mutation. Breast Cancer Res. Treat., 125, 325–49.

24. Nguyen, K., Walrafen, P., Bernard, R., Attarian, S., Chaix, C., Vovan, C., Renard, E., Dufrane, N., Pouget, J., Vannier, A., et al. (2011) Molecular combing reveals allelic combinations in facioscapulohumeral dystrophy. Ann. Neurol., 70, 627–633.

25. Cheeseman, K., Rouleau, E., Vannier, A., Thomas, A., Briaux, A., Lefol, C., Walrafen, P., Bensimon, A., Lidereau, R., Conseiller, E., et al. (2012) A diagnostic genetic test for the physical mapping of germline rearrangements in the susceptibility breast cancer genes BRCA1 and BRCA2. Hum. Mutat., 33, 998–1009.

26. Mayer, K.F.X., Rogers, J., Dole el, J., Pozniak, C., Eversole, K., Feuillet, C., Gill, B., Friebe, B., Lukaszewski, A.J., Sourdille, P., et al. (2014) A chromosome-based draft sequence of the hexaploid bread wheat (Triticum aestivum) genome. Science (80-.)., 345, 1251788–1251788.

27. Quadri, M., Vetro, A., Gismondi, V., Marabelli, M., Bertario, L., Sala, P., Varesco, L., Zuffardi, O. and Ranzani, G.N. (2015) APC rearrangements in familial adenomatous polyposis: heterogeneity of deletion lengths and breakpoint sequences underlies similar phenotypes. Fam. Cancer, 14, 41–9.

